# Probabilistic mapping of brain connections without random sampling

**DOI:** 10.64898/2026.11.12.766276

**Authors:** John Kruper, Matthew Cieslak, Ariel Rokem

## Abstract

For two decades, most tractography has mapped brain connections by sampling millions of pathways probabilistically. Sampling makes results non-deterministic and costly to store. We show that it is unnecessary: modeling tractography as an absorbing Markov chain gives the expected result of infinitely many stream-lines. This approach is more reliable than streamline-based tractography across test–retest scans. It readily maps tracts in clinical-grade data and neonates, around tumors and along small pathways. MPAT is available as open-source software at https://github.com/rokemlab/mpat

## 1 Main

Tractography based on diffusion-weighted MRI (dMRI) provides information about brain connectivity. Most computational tractography algorithms propagate three-dimensional curves, or “streamlines”, in a local greedy fashion. This approach, augmented by probabilistic algorithms and anatomical constraints, has been used to identify major white matter tracts automatically [1, 2], map whole-brain structural connectomes [3], and plan neurosurgery [4] and deep brain stimulation [5].

However, because probabilistic tractography propagates streamlines by repeated, local, stochastic steps, each run is a finite random sample from a large space of possible paths. Commonly used numbers of streamlines are insufficient to represent connectivity accurately [6], and sampling more is computationally demanding and produces large derivatives that must be stored and processed downstream.

Building on previous work that computes transition probabilities between neighboring voxels in closed form [7], we use these probabilities to instead model the white matter as an absorbing Markov chain and compute the expected result of infinitely many probabilistic streamlines without sampling. We call this method Markov Probabilistic Absorbing Tractography (MPAT) and use it to delineate the major white matter tracts with the same anatomical landmarks used in automated fiber quantification (AFQ), an existing method for recognizing tracts (using the Python implementation, pyAFQ, version 3.4 [8]).

Computing path distributions for tractography without sampling has been proposed before: both as a Fokker–Planck equation on the space of position and orientation [9] and as stochastic completion fields [10]. MPAT differs in that its transitions, propagation, and stopping rules all follow those of a standard probabilistic tractography pipeline.

Thus, we compare MPAT with probabilistic tractography in AFQ and show that probabilistic tractography approaches MPAT as more streamlines are added. In the corticospinal tract (CST) of a Human Connectome Project (HCP) participant, each tenfold increase in streamlines fills in more of the sparse regions that MPAT already captures (Fig. 1a). Even with ∼45,000 CST streamlines, from 100 million whole-brain streamlines, AFQ’s cortical endpoints remain sparser than MPAT’s (Fig. 1b). MPAT delineated the CST in ∼45 s, whereas generating 100 million streamlines took over an hour and recognizing the CST among them took a further ∼14 minutes, both multithreaded on an AMD Ryzen AI 9 HX processor. We next measure test–retest reliability in 44 subjects of the HCP test–retest dataset. MPAT was more reliable than AFQ in all 34 tracts and all 44 subjects (Dice, MPAT thresholded to match AFQ volume; mean difference +0.051; one-sided Wilcoxon signed-rank test, *p <* 0.001; Fig. 1c). In sensitivity analyses, MPAT also had higher fuzzy Dice [11] (0.819 vs. 0.772) but lower weighted Dice [2], which are nevertheless high for both algorithms (0.954 vs. 0.975). MPAT also recognizes the hippocampal cingulum and fornix using SynthSeg’s hippocampus and hypothalamus segmentations [12]. These were not reliably delineated with AFQ, but achieve a mean fuzzy Dice of 0.76 for both hippocampal cingula and 0.68–0.69 for the fornices using MPAT.

**Fig. 1.**
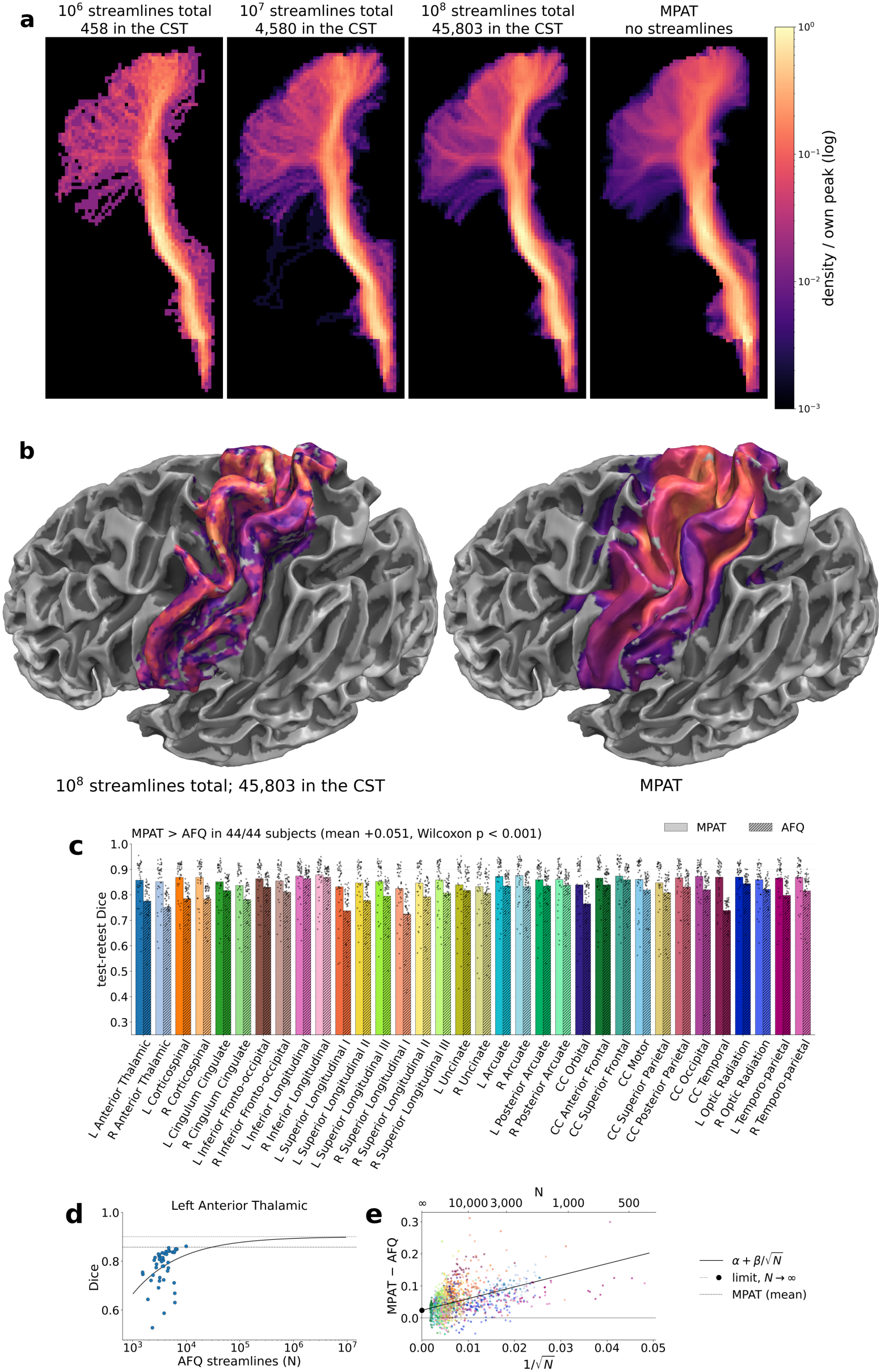
MPAT models probabilistic tractography. (a) CST (defined by SynthSeg brainstem to pre-, para- and postcentral cortex, not crossing the midline) from AFQ, subsampled to 10^6^–10^8^ whole-brain streamlines, and from MPAT. (b) Cortical CST endpoints: AFQ (left) and MPAT (right). (c) Test-retest binary Dice per tract (44 subjects; bars: mean). MPAT is higher in every subject (mean +0.051, Wilcoxon *p <*0.001). (d) AFQ’s binary Dice for the left ATR against its streamline count *N* , fitted with 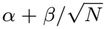 . (e) MPAT − AFQ gap for all tracts and subjects against 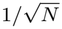. Sampling explains *∼*60% of the gap; 0.024 remains as *N* → ∞ (*p <* 0.001).

Part of this gap in reliability is expected from sampling, because a probabilistic estimate from *N* streamlines differs from its infinite-streamline limit by roughly 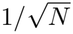 .

AFQ’s reliability rises with *N* (Fig. 1d), and a fit across subjects and tracts attributes about 60% of the gap to this term, leaving 0.024 of the difference in binary Dice as *N* →∞ (Fig. 1e). These estimates are heuristic and should be interpreted cautiously: *N* was not varied within subjects and may covary with factors, such as data quality, that also affect reliability; the 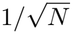 form assumes independent samples; and the remaining gap may reflect other differences between the pipelines rather than sampling.

HCP data has high spatial and angular resolution and multiple diffusion weightings. To demonstrate efficacy on an acquisition protocol that more closely resembles many research and clinical protocols, we ran MPAT on a subject from the IXI dataset (1.5 T; 15 directions of measurement and one b-value of 1,000 s/mm^2^). MPAT identifies all of the standard tracts we include in AFQ (Figure 2a). The segmentation of the left anterior thalamic radiation is atypical; however, AFQ entirely failed to recognize it in this participant, suggesting a local data-quality problem.

**Fig. 2.**
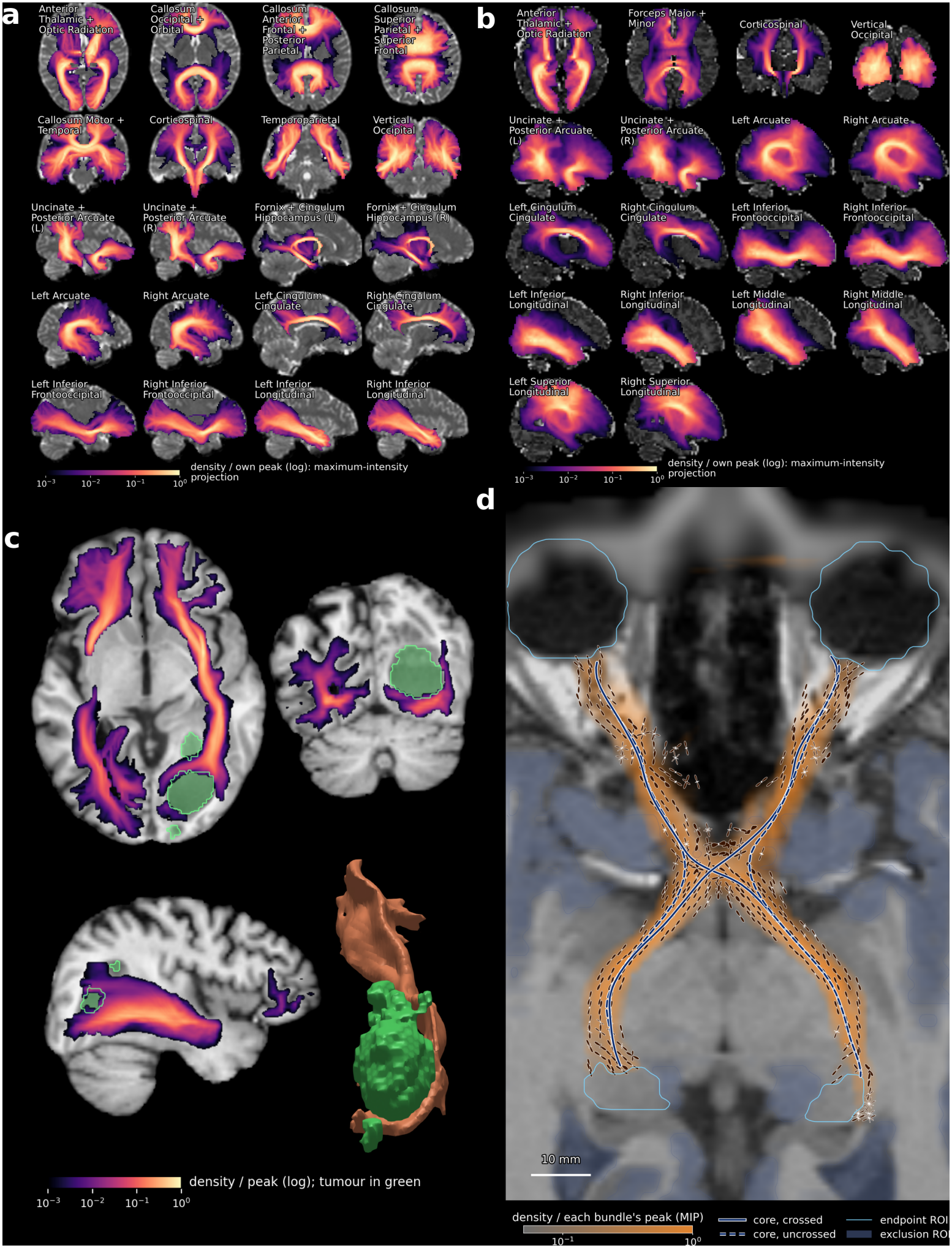
MPAT is robust. (a, b) Maximum-intensity projections of the major tracts MPAT recognizes in an IXI adult and in a neonate, respectively. (c) The inferior fronto-occipital fasciculus in a participant whose glioma has a necrotic core that displaces its course. (d) The retinogeniculate visual pathway in an HBN participant, shown on a slice tilted 25^°^ from axial.

Brain tractography and tract delineation are sensitive to atypical data and fail in situations that deviate from standard healthy brains [13, 14]. To test MPAT’s robustness, we applied it to two atypical datasets: measurements in a neonate from Grotheer et al. [13] and measurements in a brain tumor patient from the University of California San Francisco Preoperative Diffuse Glioma MRI dataset (UCSF-PDGM, #393) [15]. For neonate data, we used tissue segmentations from BabySeg [16] and the bundle definitions of BabyAFQ [13]; MPAT identified all of the BabyAFQ tracts (Fig. 2b). The UCSF-PDGM participant had a large tumor in the right occipital lobe. We treated the enhancing tumor and the edema/infiltration as white matter, so that paths can pass through them, and the necrotic core as CSF, which MPAT’s transitions never enter, so paths are steered around it rather than terminated. MPAT recovered the right inferior fronto-occipital fasciculus (IFOF) through the enhancing tumor region (Fig. 2c). In previous work, free water elimination (FWE) was needed for probabilistic tractography to track the IFOF in this setting [14]; MPAT recovered it without FWE.

MPAT can also segment small bundles that are otherwise difficult to track. Given a segmentation of the eyes, it identifies the retinogeniculate visual pathway (RGVP), including the optic nerve, chiasm, and tract, between the eyes and the lateral geniculate region of the thalamus. Typically, tracking this pathway requires specialized methods [17]. With MPAT, the RGVP emerges as the dominant pathway in a participant from the Healthy Brain Network study [18, 19], with both crossed and uncrossed fibers at the chiasm (Fig. 2d).

Because MPAT computes the expected result of tractography rather than sampling it, its outputs are differentiable: the solutions it has already computed give the gradient of any output, such as a tract profile or a cortical endpoint map, with respect to every transition probability. Errors in modeling fiber orientations could therefore be propagated into uncertainty maps alongside the density and endpoint maps, and a posterior over fiber orientations, such as the one estimated for probtrackx [20], could be carried through in the same way, producing Bayesian uncertainty on each tract without sampling a single streamline. The same solutions already give every voxel’s probability of reaching a target region, which could guide stimulation targeting: deep brain stimulation, for example, could be aimed at the subthalamic sites most connected to motor cortex and least connected to limbic regions [5].

In conclusion, by bypassing sampling, MPAT addresses many of brain tractography’s known problems [21]. It delineates the corticospinal tract in seconds rather than hours, improves test–retest reliability, and removes the need to store large trac-tograms. It is robust to atypical data and can recover small pathways. Its outputs are also differentiable, opening routes to tract-wise uncertainty maps and connectivity-based targeting. MPAT is available as open-source software at https://github.com/rokemlab/mpat.

## 2 Methods

### 2.1 MPAT

MPAT models tractography as an absorbing Markov chain. Its transient states are pairs of a white matter voxel and a heading, one of the 26 directions to the voxel’s neighbors. Its absorbing states are the voxels in which a walker ends: gray matter and the inner boundary of the brain mask (valid endings). CSF is never entered: transitions into it have zero probability, so walkers are steered around it rather than terminated, and the same holds outside the brain mask. By default, we use SynthSeg [22, 23] to generate a brain mask and gray matter, white matter, and CSF maps; voxels with white matter fraction above 0.4 inside the mask are transient, and the remaining voxels are gray matter or CSF, whichever fraction is larger.

The transition probabilities are the analytic double-ODF within-voxel transitions of Cieslak et al. [7], evaluated on fiber orientation distributions (fODFs) from constrained spherical deconvolution (CSD) [24]. We use either the single-tissue or multi-tissue [25] versions of CSD, depending on whether the data has just one non-zero b-value (single-shell) or multiple (multi-shell), respectively. Transitions are evaluated on DSI Studio’s 642-direction sphere, with steps of one voxel (of the smallest voxel side, so anisotropic voxels are handled exactly), and a maximum turning angle of 35^°^ per step. Ordering the absorbing states first, the transition matrix takes the standard form

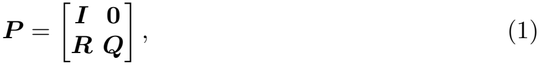

where ***I*** is the identity on the absorbing states, ***R*** holds the transitions from white matter into the absorbing states, and ***Q*** the transitions within the white matter. The fundamental matrix

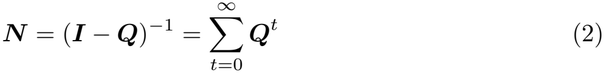

gives in *N*_*ij*_ the expected number of visits to transient state *j* of a walker started in transient state *i*. MPAT uses ***N*** in two directions. A *forward solve, x*_0_***N*** for an initial distribution *x*_0_ (a row vector over the transient states), gives the expected number of visits to every state, the occupancy, and *x*_0_***NR*** the distribution of where walkers are absorbed. A *backward solve*, ***NR*** *e* for an indicator *e* of a set of absorbing states (a column vector), gives every transient state’s probability of being absorbed in that set, the completion probability; one backward solve serves every seed. We never form ***N*** , which is dense: both solves are computed as series, 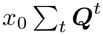 and 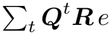, by repeated sparse products, stopped once the remaining mass falls below some tolerance (by default, 10^−6^) of the initial mass. A smaller threshold allows representation of lower-density regions.

We seed every voxel of AFQ’s white/gray matter interface that MPAT treats as white matter, with unit weight. The interface is the skeletonized inner boundary of the white matter (white matter fraction above 0.9), restricted to where gray matter outweighs CSF [8]. Each seed’s weight is divided equally among the voxel’s fODF peaks, found with DIPY’s peak finder (relative threshold 0.25, minimum separation 45^°^) [26]; each peak launches a pair of walkers in opposite directions along it, the two halves of one streamline, mimicking a seeding strategy common in probabilistic tractography. Summed over headings, *x*_0_***N*** is then the streamline density of each voxel (in expected walker steps), and *x*_0_***NR*** the streamline endpoint density.

### 2.2 MPAT Tract Recognition

A streamline belongs to a tract when its two halves together satisfy the tract’s ROIs: it avoids every exclusion ROI, passes through every inclusion ROI, and ends validly (gray matter or the edge of the brain) with one end in each endpoint ROI. MPAT implements all three of these types of ROIs:

- *Exclusion ROIs*: With ***M*** the diagonal mask that is 0 on excluded states and 1 elsewhere,

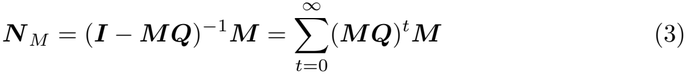

is the fundamental matrix of the chain in which excluded states are invalid endings: it counts only visits made before a walker reaches an excluded state.
- *Endpoint ROIs*: Let *e*_*S*_ and *e*_*E*_ indicate the valid absorbing states in the start and end ROIs. The backward solves

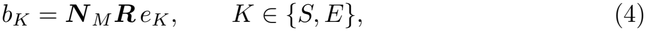

give every transient state’s probability of finishing in *K*. The two halves of a streamline walk independently, so a half contributes only as often as its partner finishes at the other end. Weighting each half’s launch distribution *a*_*h*_ by its partner’s completion,

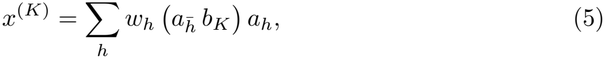

(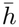 the partner of *h, w*_*h*_ the seed weight) and walking forward gives the bundle’s density as the product of occupancy and completion,

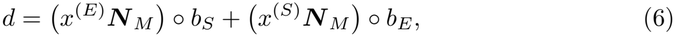

the visits of walkers that will go on to finish in *S* (or *E*) while their partners finish in *E* (or *S*). This product of a forward and a backward solution is the stochastic completion field of Williams and Jacobs [27] and the reactive density of transition path theory [28]. Bundles without endpoint ROIs use the indicator of every valid ending in place of *e*_*S*_ and *e*_*E*_.
- *Inclusion ROIs*: Passing through an ROI is the complement of avoiding it, and avoiding is an exclusion. For inclusion ROIs *A*_1_, … , *A*_*m*_, inclusion–exclusion gives

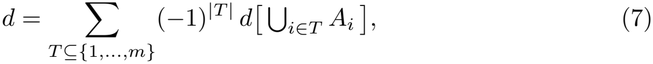

where *d*[*X*] is Equation 6 with *X* added to the exclusion ROIs. Writing *X*_*T*_ = *X* ⋃U_*i* ∈*T*_ *A*_*i*_ for the exclusion ROIs *X* together with the inclusion ROIs in *T* , and subscripting Equations 3–5 by the exclusion set they use, the general equation for the density of a tract, including all types of ROIs, is

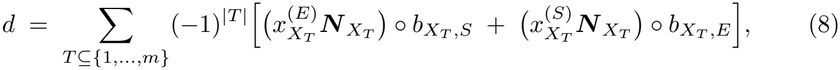

with *b*_*X,K*_ = ***N*** _*X*_ ***R*** *e*_*K*_ and 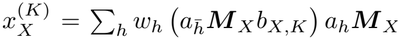. Note there are 2^*m*^ terms, and each term costs one backward and one forward solve, so inclusion ROIs are significantly more expensive to add than the other two types.

### 2.3 Tract definitions

MPAT uses AFQ’s tract definitions [1, 2], with the following additions. Tracts that do not cross the midline are kept to their hemisphere exactly, voxel by voxel. In the corticospinal tracts, a tolerance of 10 mm past the midline is allowed, because the tracts run next to the midline in the brainstem, and an exact cut can sever them in slightly misaligned data. The anterior thalamic radiations exclude a box on the midline, beneath the corpus callosum and above the thalamus, that contains the fornix and septum pellucidum (MNI *x* ∈ [−3, 3], *y* ∈ [−28, 15], *z* ∈ [−5, 22] mm), as walkers otherwise reach the thalamus along the fornix. In addition, three new tract definitions are used:

#### Corticospinal tract (Fig. 1a,b)

For the comparison with probabilistic tractography, we defined each corticospinal tract by its endpoints alone, without waypoints: SynthSeg’s precentral, paracentral and postcentral cortex [22] and SynthSeg’s brainstem, kept to its hemisphere as above. These labels come from the subject’s own segmentation, so they were used as they are, without the 4 mm dilation AFQ applies to endpoint ROIs registered from a template, in both MPAT and AFQ.

#### Hippocampal cingulum

The two waypoint ROIs of AFQ’s hippocampal cingulum [1], with one end in SynthSeg’s hippocampus.

#### Fornix

From SynthSeg’s hippocampus to SynthSeg’s hypothalamus [12], through a waypoint on the fornix body (MNI *x* ∈ [−4, 4], *y* ∈ [−25, −5], *z* ∈ [−8, 22] mm, warped to the subject), kept to its hemisphere within 3 mm of the midline. The waypoint requires the long route along the crus and body, over the thalamus, rather than the direct route beneath it.

### 2.4 Comparison with AFQ, reliability, and statistics

We compared MPAT with AFQ implemented in Python (pyAFQ 3.4) [2, 8]. For the test–retest analysis (Fig. 1c–e), AFQ was run on each session with its default trac-tography and tract recognition, using the same multi-shell multi-tissue CSD MPAT used [25], with SynthSeg’s brain mask and tissue segmentation. Each tract’s stream-lines were converted to a density map. The vertical occipital fasciculus was left out of the comparison because the current version of pyAFQ implements additional steps for division into sub-bundles (unpublished), which are not yet implemented in MPAT.

For the convergence comparison (Fig. 1a,b), we ran AFQ’s probabilistic tractography from 5 ×10^7^ random seeds in HCP subject 627549, producing 1.2 ×10^8^ whole-brain streamlines, and recognized the left corticospinal tract as defined above (minimum length 40 mm, without AFQ’s outlier cleaning). To show the tract at 10^6^, 10^7^ and 10^8^ whole-brain streamlines, we drew random subsets of the recognized streamlines, without replacement, in proportion to each target count.

Test–retest reliability was measured with three overlap measures, each computed between the two sessions’ density maps of a tract. The primary measure was the binary Dice coefficient, 2 |*A* ∩ *B /*(|*A*| + |*B*|), on masks matched in volume. AFQ’s mask in each session was every voxel reached by at least one streamline, and *K* was the mean of the two sessions’ mask sizes. MPAT’s mask in each session was its *K* densest voxels. As sensitivity analyses, we computed the fuzzy Dice coefficient of Crum et al. [11], FD = ∑_*v*_ min(*p*_*v*_, *q*_*v*_), where *p*_*v*_ and *q*_*v*_ are the tract’s densities at voxel *v* in the two sessions after each map has been normalized to sum to one; FD is 1 for identical maps and requires no threshold. We also computed the weighted Dice coefficient [29] used by Kruper et al. [2],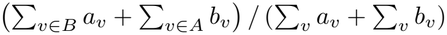, where *a* and *b* are the two sessions’ density maps, and *A* and *B* are the same volume-matched masks used for the binary Dice coefficient. Weighted Dice weights each voxel by its density, so it measures how much of each map’s density falls inside the other session’s mask, not how well the densities agree. We compared the 34 tracts both methods recognize, excluding the vertical occipital fasciculus, which AFQ subdivides. AFQ recognized all 34 tracts in both sessions of every subject. Each subject’s binary Dice score was averaged over these tracts, and MPAT and AFQ were compared across subjects with a one-sided Wilcoxon signed-rank test.

To estimate the contribution of sampling, we took *N* as the harmonic mean of AFQ’s streamline counts for a tract in the two sessions. For each tract we fitted AFQ’s binary Dice as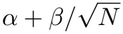(Fig.1d), and across tracts and subjects we fitted the gap between the methods as 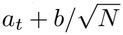, with a separate intercept *a*_*t*_ for each tract and a shared slope *b*, by least squares within tracts (Fig. 1e). The gap remaining as *N* → ∞ is the mean of the *a*_*t*_; its 95% confidence interval and one-sided *p*-value come from 5,000 bootstrap resamples of subjects, with each subject’s tracts resampled together.

### 2.5 HCP

We analyzed HCP test-retest data from 44 neurologically healthy participants between 22 and 35 years of age. Full details of the HCP acquisition protocol are given in [30]. In short, diffusion-weighted data were collected on three shells (b ≈ 1,000 s/mm^2^, b ≈2,000 s/mm^2^, and b ≈3,000 s/mm^2^), each using a TR of 5.5 s and a TE of 89 ms. For every shell, 90 non-collinear directions were acquired twice, once with each of two inverse phase-encoding directions (LR and RL), and five non-diffusion-weighted acquisitions were selected, giving 190 data points per shell in total [31]. Data had a spatial resolution of 1.25 × 1.25 × 1.25 mm^3^. Acquisition was performed with multislice echo planar imaging and multiband excitation on a 3T Siemens Skyra scanner customized for diffusion imaging, equipped with an SC72 gradient coil and an upgraded gradient power supply (Gmax = 100 mT/m; Gmax = 97.4 mT/m in practice after optimization for gradient duty cycle). Signal was received in parallel through a 32-channel head coil.

The data we used were supplied by HCP after processing with the HCP minimal preprocessing pipelines described in [32]. In short, intensity was normalized across the six acquisition series using the non-diffusion-weighted (b0) images. Pairs of b0 images with opposing phase encoding were then used to estimate and correct EPI distortions with the FSL “topup” tool [33]. The FSL “eddy” tool [34] was used to correct artifacts caused by eddy currents and motion, and gradient spatial non-linearities were computed [35]. The spatial transform between the average b0 image and the T1-weighted data was calculated with a boundary-based registration approach (FSL FLIRT BBR and FreeSurfer’s “BBRegister” [36]). Lastly, the eddy-corrected data were transformed into the 1.25 mm structural volume space in one step that applied both the gradient nonlinearity correction and the T1w registration.

### 2.6 IXI

We used a single participant from the Information eXtraction from Images (IXI) dataset. This participant’s data was acquired on a 1.5T Philips Medical Systems Gyroscan Intera scanner at a spatial resolution of 1.75 × 1.75 × 2.35 mm. Diffusion-weighted images were collected along 15 directions, along with one b0 image (TR = 9054 ms, TE = 80 ms, 3 averages). The T1-weighted image was acquired with TR = 9.8 ms, TE = 4.6 ms, and a flip angle of 8^°^. We preprocessed the data with QSIPrep 26.1.1 using the default settings and an output resolution of 1.75 mm.

### 2.7 Neonate Data

We used data from a single healthy, full-term neonate from Grotheer et al. [13], which was scanned during natural sleep on a 3T GE Discovery MR750 scanner with a 32-channel head coil. Multi-shell dMRI was acquired at 2 mm isotropic resolution (9 b = 0 volumes, 30 directions at b = 700 s/mm^2^, and 64 directions at b = 2000 s/mm^2^; TE = 75.7 ms, TR = 2800 ms), along with a reverse phase-encoding b = 0 scan. Preprocessing followed Grotheer et al. [13] and included PCA denoising, FSL topup and eddy for distortion, eddy-current and motion correction, and ANTs bias correction. As described above, tissue segmentations were generated with BabySeg [16].

### 2.8 UCSF-PDGM

The UCSF-PDGM dataset is a public collection of preoperative imaging and clinical data from adult patients with diffuse gliomas [15]. It includes T1 and T2 MRI, DTI, and segmentations of the tumor components (enhancing tumor, necrotic core, and edema/infiltration regions). The protocols have been described previously. In short, diffusion-weighted MRI was acquired at 2 mm isotropic resolution with 55 directions and b = 2,000 s/mm^2^.

### 2.9 HBN

We randomly selected 1 subject from the HBN pediatric mental health study to evaluate the RGVP [18]. HBN includes dMRI data from 1651 participants between 5 and 21 years of age. Full descriptions of the acquisition and preprocessing procedures are given elsewhere [18, 19]. In short, data were collected on 3T Siemens scanners at three sites in the New York area, with a spatial resolution of 1.8 × 1.8 × 1.8 mm. Two shells, at b ≈1000 s/mm^2^ and b ≈2000 s/mm^2^, were each measured with 64 non-collinear directions. Preprocessing was performed using QSIPrep as described previously [19, 37].

Brain masks rarely include the eyes, so we defined the tracking domain as the convex hull of the thalamus and both eyes, and treated every voxel within it as trackable, not only those SynthSeg labels as white matter. SynthSeg [22] provided labels for the thalamus and cortex. All non-thalamic gray matter was an exclusion ROI. The end-point ROIs were the eyes and the lateral, posterior, inferior eighth of each thalamus, as an approximation of the lateral geniculate nucleus; the remainder of the thalamus was an exclusion ROI. The thalamic endpoint accepted paths only from its nine lateral neighboring voxels.

Each eye was segmented by warping an approximate eye center from MNI space (left: [−30.5, 56.4, −39.7]; right: [29.3, 56.4, −40.3]) into the subject with AFQ’s registration, then fitting a sphere (radius 9–13 mm) to the arc of the globe’s back wall on the unmasked T1-weighted image. The sphere was found with a gradient-directed Hough transform followed by a least-squares refit, so it needs only part of the wall and works on face-blurred or defaced images.

## Acknowledgements

John Kruper was supported through the Weill NeuroHub at the University of Washington. This research was enabled in part by support provided by the Digital Research Alliance of Canada. Data were provided in part by the Human Connectome Project, WU-Minn Consortium (Principal Investigators: David Van Essen and Kamil Ugurbil; 1U54MH091657) funded by the 16 NIH Institutes and Centers that support the NIH Blueprint for Neuroscience Research; and by the McDonnell Center for Systems Neuroscience at Washington University. IXI data were obtained from the Information eXtraction from Images project (https://brain-development.org/ixi-dataset/). This manuscript was prepared using data obtained from the Child Mind Institute Biobank, Healthy Brain Network dataset, and reflects the views of the authors and does not necessarily reflect the opinions or views of the Child Mind Institute. UCSF-PDGM data were obtained from The Cancer Imaging Archive (TCIA) [15, 38, 39].

## Code availability

MPAT is available at https://github.com/rokemlab/mpat. Scripts to reproduce figures are available at https://github.com/36000/mpat_paper/.

## Data availability

All data are publicly available. HCP data are available from the Human Connectome Project (https://db.humanconnectome.org) under the WU-Minn HCP data use terms. IXI data are available from https://brain-development.org/ixi-dataset/. The neonate data are from Grotheer et al. [13], available at https://figshare.com/ndownloader/files/38053692. UCSF-PDGM data are available from The Cancer Imaging Archive [15, 38]. HBN data are available from the Child Mind Institute (https://fcon_1000.projects.nitrc.org/indi/cmi_healthy_brain_network/) under its data use agreement.

## Ethics declarations

This study used only de-identified data from public datasets, each collected with approval from the relevant institutional review boards.

## Author contribution

JK: Conceptualization, Methodology, Software, Analysis, Writing - Original Draft. Writing - Review & Editing. MC: Conceptualization, Methodology, Software, Writ- ing - Review & Editing. AR: Conceptualization, Supervision, Project administration, Funding acquisition, Writing - Original Draft. Writing - Review & Editing.

## Competing interests

The authors declare no conflicting interests

## Notes

https://github.com/rokemlab/MPAT

https://github.com/36000/mpat_paper

### Competing Interest Statement

The authors have declared no competing interest.

